# The Dual Mechanisms of Cognitive Control (DMCC) Project

**DOI:** 10.1101/2020.09.18.304402

**Authors:** Todd S. Braver, Alexander Kizhner, Rongxiang Tang, Michael C. Freund, Joset A. Etzel

## Abstract

The Dual Mechanisms of Cognitive Control (DMCC) project provides an ambitious and rigorous empirical test of a theoretical framework that posits two key cognitive control modes: proactive and reactive. The framework’s central tenets are that proactive and reactive control reflect domain-general dimensions of individual variation, with distinctive neural signatures, involving lateral prefrontal cortex (PFC) in interactions with other brain networks and circuits (e.g., frontoparietal, cingulo-opercular). In the DMCC project, each participant is scanned while performing theoretically-targeted variants of multiple well-established cognitive control tasks (Stroop, Cued Task-Switching, AX-CPT, Sternberg Working Memory) in three separate imaging sessions, that each encourage utilization of different control modes, plus also completes an extensive out-of-scanner individual differences battery. Additional key features of the project include a high spatio-temporal resolution (multiband) acquisition protocol, and a sample that includes a substantial subset of monozygotic twin pairs and participants recruited from the Human Connectome Project. Although data collection is still continuing (target N=200), we provide an overview of the study design and protocol, planned analytic approaches and methodological development, along with initial results (N=80) revealing novel evidence of a domain-general neural signature of reactive control. In the interests of scientific community building, the dataset will be made public at project completion, so it can serve as a valuable resource.

## Introduction

The DMCC project is a large-scale longitudinal study that provides a systematic test of the Dual Mechanisms of Control (DMC) framework, a theoretical account of both inter- and intra-individual variability in cognitive control mechanisms (Braver, 2012; Braver et al., 2007). The study has multiple components, which assess brain activity and behavioral performance via a newly developed task battery. This battery is designed to probe the two main control modes postulated in the framework, proactive and reactive, in terms of both experimental manipulations and individual differences. The sample size target of the DMCC project is to acquire data from 200 healthy young adults. It is worth noting that although there have now been a number of larger-scope neuroimaging projects that share some similarity with the DMCC (e.g., HCP, ABCD, Cam-Can, IMAGEN, PNC, UK Biobank; Casey et al., 2018; Essen et al., 2013; Miller et al., 2016; Satterthwaite et al., 2014; Schumann et al., 2010; Shafto et al., 2014), this project is somewhat unique in that was designed to test a specific theoretical framework, and with sufficient sample size for not only robust and reliable estimation of group effects, but also to provide sensitivity to individual differences effects, though subject to the constraints of what is financially and logistically feasible within a single-lab, single-PI effort. Likewise, the sample also has unique features, consisting of two informative participant subsets: 1) identical (monozygotic) twin-pairs, enabling phenotypic analyses of genetic and/or environmental similarity effects; and 2) participants recruited from the Human Connectome Project (HCP), enabling integration of DMCC data with prior HCP data.

Funded by a NIMH MERIT award, primary data collection in the DMCC project began in late 2016. As of 2020, the project is still on-going (thanks to a second round of NIMH funding), with data collection expected to continue until 2023. Each participant takes part in at least one wave of testing that an extensive out-of-scanner behavioral session and three fMRI neuroimaging sessions. In the neuroimaging sessions, high-resolution anatomical, resting state fMRI, and task fMRI data are collected. In total, a minimum of 300 minutes (5 hours) task-fMRI and 30 minutes of resting-state fMRI are collected for each participant. The out-of-scanner assessments include over 25 measures of cognitive ability and personality traits, as well as psychological and physiological indices of health and well-being. In each fMRI session, participants perform the full DMCC task battery, in one of three conditions, baseline, proactive and reactive, with these conditions describing distinct variants of each task in the battery. The purpose of this report is to describe the origin and current state of the DMCC project, including the underlying theoretical and conceptual backbone behind the DMCC protocol, the task paradigms that comprise the primary neuroimaging battery, and the analysis pipeline, including some aspects of current methodological development. Most importantly, we present a first set of initial results from the project, which provide new evidence of a consistent neural signature of reactive control.

## Theoretical and Conceptual Goals of the DMCC Project

The DMCC project was explicitly designed to help achieve the goals of the Research Domain Criteria (RDoC), a major NIMH strategic initiative. The RDoC initiative aims for a reconceptualization of mental illness and neuropsychiatric disorders in terms of underlying dimensions that can be characterized at different levels of analysis, from behavioral profiles, to neural system and circuit abnormalities, all the way to genetic and molecular causes (Cuthbert & Insel, 2013). The DMCC project was structured to help achieve these goals, by providing *a rigorous and systematic examination of the putative core dimensions and neural mechanisms that give rise to variation in cognitive control function*.

Cognitive control, which refers to the ability to regulate, coordinate, and sequence thoughts and actions in accordance with internally maintained goals (E. K. Miller & Cohen, 2001), is one of the key domains or constructs of focus within the RDoC initiative. There is a strong consensus that cognitive control impairments are a critical component of a wide-range of mental health and neuropsychiatric disorders (e.g., schizophrenia, depression, ADHD, Parkinson’s, Alzheimers). Yet, there is still a poor understanding of the underlying sub-components and mechanisms that give rise to both normal and pathological variation in cognitive control function. In developing the DMCC project, we relied heavily on the DMC theoretical framework, which we believe provides critical experimental leverage and methodological tools for uncovering and characterizing the component mechanisms of cognitive control variation.

The DMC framework is a unifying and coherent theoretical account that explains three empirically observed sources of variation – within-individual (task and state-related), between-individual (trait-related), and between-groups (i.e., impaired populations with changes to brain function and integrity) – in terms of an underlying core dimension of variability related to the temporal dynamics of cognitive control. It is this emphasis on cognitive control variability and temporal dynamics that critically distinguishes the DMC framework from other theoretical accounts, which nonetheless posit similar computational mechanisms and neural architectures (Banich, 2009; Botvinick et al., 2001; Engle & Kane, 2003; Herd et al., 2014; Koechlin & Summerfield, 2007; E. K. Miller & Cohen, 2001; Miyake et al., 2000)

In the DMC framework, the key distinction is between two control modes that have contrasting dynamic neural signatures. Proactive control involves sustained and preparatory activation of cognitive goal representations within lateral prefrontal cortex (PFC), enabled by phasic inputs (for goal updating) and tonic signals (for goal maintenance) arising from the mid-brain dopamine (DA) system (and associated components, i.e., dorsal and ventral striatum). In contrast, reactive control involves transient, stimulus-driven goal activation in these lateral PFC regions, based on signals arising from neural circuits that mediate interference/conflict detection (e.g., anterior cingulate cortex, medial frontal cortex; ACC/MFC) and/or episodic/associative cueing (e.g., posterior parietal cortex [PPC], medial temporal lobe [MTL]).

Based on a range of theoretical arguments (detailed in Braver, 2012; Braver et al., 2007), we postulate that the proactive and reactive control modes reflect computational tradeoffs with complementary costs and benefits. Consequently, successful cognition depends upon a variable mixture of proactive and reactive control strategies. Moreover, there are a variety of state, trait, and population factors that influence which control mode is dominant, including available cognitive resources and capacity, motivational salience of task performance and reward attainment, expectations for interference, and the integrity/efficacy of relevant neural systems and circuits.

A key tenet of the DMC framework is that proactive and reactive control appear to serve as meaningful constructs in the RDoC sense, in that they appear to: a) index coherent dimensions of both state and trait-related variability in normal cognitive control function; b) be useful for characterizing both age-related changes and clinical impairment in a variety of populations; and c) exhibit unique and well-defined behavioral and neural signatures. Indeed, a tantalizing possibility is that proactive and reactive control might act as endophenotypes, in also reflecting meaningful dimensions of genetic variation (e.g., single nucleotide polymorphisms [SNPs], such as COMT; Furman et al., 2020; Green et al., 2012; Mier et al., 2009). For example, we previously speculated that the complementary computational tradeoffs of proactive and reactive control could each confer evolutionary advantages optimized for different environmental contexts (e.g., stable / predictable vs. rapidly changing / chaotic), leading to their stable expression in the population (Braver et al., 2010).

A major aim of the DMCC project is to extend our understanding of proactive and reactive control, by conducting a comprehensive test of their construct validity. Although rigorous establishment of construct validity is a critically important endeavor (Cronbach & Meehl, 1955), particularly with respect to RDoC goals, it is still only infrequently attempted in investigations of cognitive (executive) control (Friedman & Miyake, 2016; Karr et al., 2018; Rey-Mermet et al., 2019), and is even more rarely a focus of cognitive neuroscience research in this domain (Derrfuss et al., 2004; Kragel et al., 2018; Sylvester et al., 2003). A first step is to establish convergent validity, which requires assessment of multiple distinct measures of proactive and reactive control in a within-subject design, in order to test for common cross-task relationships and patterns of activation. A second step is to establish divergent (discriminant) validity, by demonstrating that proactive and reactive control do in fact reflect dissociable constructs. This is actually quite challenging experimentally, in that proactive and reactive control are by definition temporally related, such that reduced utilization of proactive control will increase the demand on reactive control, and vice versa (which we have previously termed a *reactive-proactive shift;* Braver et al., 2009). The third step of construct validation is to properly situate proactive and reactive control within a nomological network of related constructs at different levels of mechanism and description, including other measures of individual difference (e.g., personality/motivation, intelligence, working memory capacity), brain function (e.g., anatomy, connectivity), and genetics (e.g., heritability, SNPs). Finally, for the longer-term effort of evaluating the utility of proactive and reactive control as meaningful constructs in studies of health and disease, it will be necessary to determine their predictive validity for important functional outcomes (e.g., educational and career achievement, physical and mental-health status and vulnerabilities).

In this first stage of reporting on the project and its progress, we focus on the new cognitive control battery developed to provide dissociable measures of proactive and reactive control that are also convergent across multiple task paradigms. Additionally, we detail our approach to data acquisition and analysis, while also describing current methodological developments, both of which are aimed at achieving both high degrees of reproducibility, transparency, and ensuring quality control / quality assurance (QC/QA).

## The DMCC Task Battery

The DMCC task battery includes four well-established task paradigms frequently used in the cognitive control literature: Stroop, AX-CPT, Cued Task-Switching and Sternberg Working Memory. Critically, however, each of the tasks is performed under three different conditions that encourage utilization of different cognitive control strategies: baseline, proactive, and reactive. Moreover, the variants of these paradigms adopted within the DMCC project are in some cases novel, without prior precedent in the literature. Consequently, here we provide an overview of each of the tasks in the battery, along with the rationale for their inclusion and the logic behind the different manipulations (see Figure 1). It is also worth noting that the full DMCC task battery is being explored in a parallel behavioral study (conducted on-line through Amazon MTurk), via a test-retest format, to explore psychometric properties. A report of behavioral findings from that study is forthcoming, so here we emphasize the basic behavioral effects, along with predictions for neuroimaging.

**Figure 1.**
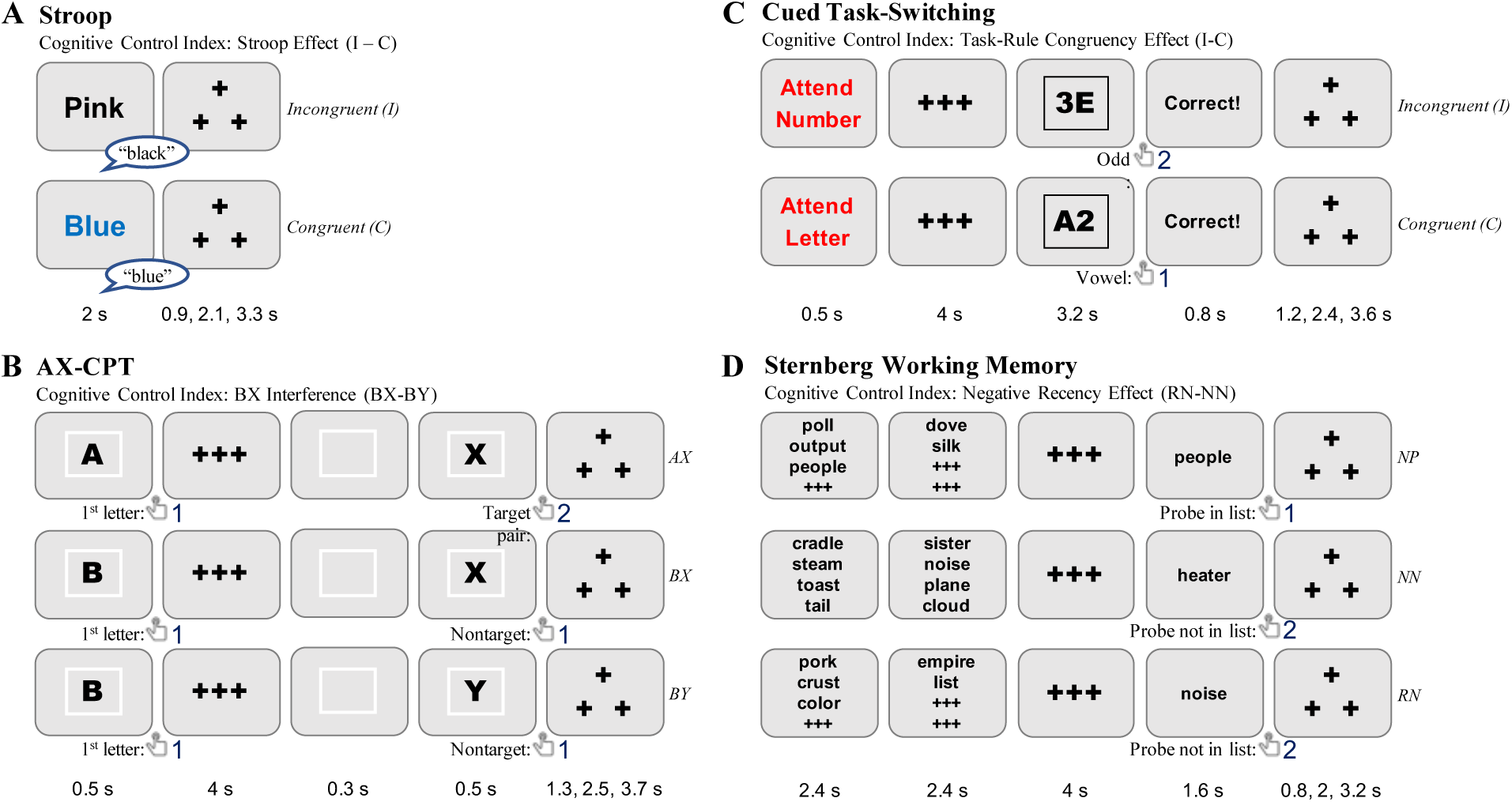
Diagram of cognitive control tasks used in DMCC battery, highlighting key trial contrasts used to define cognitive control index. A. Stroop task, illustrating contrast between incongruent and congruent trials. B. AX-CPT, illustrating target (AX) trials as well as key contrast between BX and BY trials. C. Cued-Task Switching, illustrating contrast between incongruent and congruent trials. D. Sternberg Working Memory, illustrating target (NP) trials, as well as key contrast between RN and NN trials. Diagram illustrates Baseline condition, with stimulus timing information listed for each task.

### Stroop

The color-word Stroop is widely recognized as a canonical task of cognitive control, in which top-down selective attention is required to focus processing on the task-relevant font color of printed words, while ignoring the irrelevant but otherwise dominant word name. The primary index of cognitive control is thus the *Stroop interference effect*, which contrasts incongruent (word name indicates a different color than the font color, e.g., BLUE in red font) and congruent (word name matches font color, e.g., BLUE in blue font) trials (see Figure 1A). A key dimension of the task that has often been used to manipulate cognitive control demands is that of probability congruence (PC; Bugg et al., 2008; Bugg & Crump, 2012). Under high PC conditions, congruent trials are frequent and incongruent trials are rare, such that cognitive control demands are on average low and intermittent. Consequently, the *baseline* condition utilizes high PC, as it produces robust Stroop interference and individual differences effects (Kane & Engle, 2003).

In contrast, the *proactive* condition utilizes a low PC condition in which PC is decreased in a global, or list-wide manner (Bugg, 2012; Bugg et al., 2011). In this case, proactive control is theoretically associated with sustained maintenance of the task goal to attend to the ink color dimension and ignore the word, which should be present in a consistent (i.e., on all trials) and persistent manner (i.e., engaged even prior to stimulus onset). Thus, the key prediction is that Stroop interference should be reduced on all trials, relative to baseline condition with high PC conditions. Likewise, in terms neural activity dynamics, the key prediction is that cognitive control-related mechanisms should be engaged prior to stimulus onset, potentially sustained across trials, and/or present globally on all trials.

The *reactive* condition also manipulates PC, but in an item-specific, rather than list-wide fashion. In this case, specific colors occur with low PC (e.g., RED in green font is frequent, while RED in red font is rare), while others occur with high PC (e.g., YELLOW in yellow font is frequent). This type of item-specific PC manipulation is theoretically predicted to enhance the utilization of *reactive control* when low PC items are encountered. For these items, strong associations may develop between a critical feature (a specific ink color) and increased control demands (i.e., high interference), leading to this feature more effectively engaging goal retrieval and utilization (Bugg et al., 2011; Bugg & Hutchison, 2013; Jacoby et al., 2003). When this occurs, the engagement of cognitive control-related neural activity would be expected to be transient, present only after stimulus onset, and primarily engaged by low PC incongruent items (relative to low PC congruent items).

A novel feature of the Stroop tasks included in the battery, is that there are actually two distinct, and intermixed set of items, biased items, for which PC is manipulated across conditions, and unbiased items (“PC-50”: 50% congruent, 50% incongruent). The unbiased items, which are included and equivalent across all three conditions, enable tighter comparisons and predicted dissociations among conditions. Specifically, in the *proactive condition*, changes in Stroop behavioral interference and neural activity, relative to baseline, should equivalently impact both types of items, whereas in the *reactive condition*, the cognitive control-related changes should be specific to the biased items. Finally, it is worth noting that because of the large numbers of different font colors (8) included in each of the conditions, the task is implemented with vocal rather than manual responding, using digitization and automated signal processing algorithms to extract response latencies from the noisy scanner environment. In our previous behavioral studies, we have used similar experimental manipulations to dissociate proactive and reactive control, using both picture-word (Gonthier et al., 2016), and color-word (Gourley et al., 2016) Stroop variants.

### AX-CPT

The AX-CPT has become increasingly utilized as a task of context processing and cognitive control, given its simplicity, flexibility and applicability in a wide-range of populations (Barch et al., 2008; Chatham et al., 2009; Chun et al., 2018; MacDonald, 2008; Paxton et al., 2007). In the task, contextual cues constrain the appropriate response to probe items. As the name suggests, A-cues followed by X-probes require a target response, and occur with high frequency, leading to strong cue-probe associations. Cognitive control is postulated as key process involved in maintaining and utilizing the contextual cue information, in order to minimize errors and response interference occurring on BX trials, which occur when the X-probe is presented but not preceded by an A-cue. Thus, a useful index of cognitive control for this task is the *BX interference effect*, which contrasts BX and BY (neither an A-cue nor an X-probe is presented, leading to low control demands) trial types (see Figure 1B). In prior work, shifts in the tendency to utilize proactive or reactive control have not only been observed when comparing different populations or groups, but have also been manipulated within-subjects (Braver et al., 2009).

The AX-CPT conditions included in the battery extend prior recent work, by using a task variant in which the A- and B-type contextual cues occur with equal frequency, thus eliminating confounds in earlier versions that could be due to the lower overall frequency of encountering B-cues (Gonthier, Macnamara, et al., 2016; Richmond et al., 2015). Further, these conditions also include no-go trials, in which the probe is a digit rather than letter. Because of the increase in response uncertainty (i.e., three types of probe response are possible: target, nontarget, no-go), the addition of no-go trials decreases the overall predictive utility of context information for responding, and as a consequence was found to reduce the overall proactive control bias typically observed in healthy young adults. Thus, the *baseline* condition includes these no-go trials to produce a “low control” state, from which to more sensitively observe condition-related changes in control mode (Gonthier, Macnamara, et al., 2016).

The *proactive* condition replicates prior work using context strategy training to increase predictive preparation of responses following contextual cue information (Edwards et al., 2010; Gonthier, Macnamara, et al., 2016; Paxton et al., 2006). Specifically, prior to performing this condition, participants are provided with explicit information regarding the frequencies of these cue-response associations, and receive training and practice in utilizing them to prepare the dominant responses. In addition, during inter-trial intervals, participants are provided with visual instructions to “remember to use the strategy”. The key prediction is that the increased utilization of contextual cue information will lead to a bias to prepare a target response following an A-cue (analyzed in terms of both AX and AY trials) and a nontarget response following a B-cue, leading to reduced interference on BX trials, but a side effect of which will be increased interference on AY trials, which occur when the A-cue is *not* followed by an X-probe. This translates into a prediction of increased cue-related neural activity, which might also be accompanied by sustained activation (in maintaining the instructed strategy) across the task block.

The *reactive* condition utilizes a new manipulation which has not previously been examined in prior work. Specifically, in the reactive condition item-specific probe cueing is present (similar to other cueing manipulations in tasks, such as the flankers; Braem et al., 2019; Bugg & Crump, 2012; Crump et al., 2006), such that on high control demand trials (AY, BX, nogo) the probe item appears in a distinct spatial location, and with a distinct border color surrounding it (presented briefly before the onset of the probe). Critically, because these stimulus-control associations (i.e., between border color / spatial location and high control demand) only form at the time of probe onset, they are not hypothesized to modulate the utilization of proactive control strategies. Likewise, the probe features are insufficient to direct stimulus-response learning, since they do not directly indicate the appropriate response to be made (i.e., either a non-target or no-go response could be required). In contrast, because probe features serve as cues signaling high control demand, and they can drive more rapid and effective retrieval of contextual information to resolve the conflict. The key prediction is that utilization of probe features should reduce BX interference in the reactive condition. In terms of neural activity dynamics, reactive control effects should be observable as transient probe-triggered activation on BX (relative to BY trials).

### Cued-TS

Cued task-switching (Cued-TS) has long been recognized as a critical paradigm to assess a core component of cognitive control – the ability to update and activate task-representations in on-line manner in order to appropriately configure attention and action systems for processing the upcoming target (Kiesel et al., 2010; Meiran, 1996; Vandierendonck et al., 2010). The key aspect of the paradigm is that two or more tasks randomly alternate in a trial-by-trial fashion, with target items typically being ambiguous, so that they can be processed according to multiple task rules. Consequently, the advance presentation of the task cue, prior to target onset, is what disambiguates the target and specifies the appropriate stimulus-response rules. An important index of cognitive control in task-switching paradigms is the *task-rule congruency effect (TRCE;*Meiran & Kessler, 2008), which refers to the increased interference (both errors and reaction time) when the target response required for the current task trial is incongruent to the response that would be required to the same target stimulus if a different task had been cued (see Figure 1C). For example, in letter-digit task-switching (also called consonant-vowel, odd-even or CVOE; Minear & Shah, 2008; Rogers & Monsell, 1995), if in the letter task, a right button press is required for a consonant, and left button press for a vowel, but in the digit task, a right button press is required for odd and left button press for even, the “3E” target stimulus would be incongruent (whereas the “A2” target stimulus would be congruent, since for either task, the left button press would be correct). Another cognitive control effect is the mixing cost, which occurs on all trials in task-switching blocks (relative to single-task blocks; Rubin & Meiran, 2005))

In prior work, it was found including reward incentives on a subset of trials, with reward cues presented at the time of the task cue, led to a strong reduction in the mixing cost (interference on task-repeat vs. single-task trials) – and this was present even on the trials that were non-incentivized – but no effect on the TRCE (Bugg & Braver, 2015) This finding was interpreted as indicating that global performance enhancements are associated with *proactive control*, whereas *reactive control* primarily influences the TRCE, and so is less impacted by advance reward incentive manipulations. The Cued-TS conditions included in the DMCC battery build on these prior findings, by using variants of the CVOE (letter/digit) paradigm that accentuate the robustness of the TRCE, as a putative marker of *reactive control*, while also incorporating advance task cues with a long cue-to-target interval, which enables effective utilization of *proactive control*.

In the *baseline* condition, target stimuli are list-wide mostly congruent, as prior work has found that mostly congruent conditions result in a large and robust TRCE (Bugg & Braver, 2015). The *proactive* condition follows Bugg & Braver (2015) in keeping the same list-wide mostly congruent structure as the baseline condition, but adding reward incentives on a subset of trials. Specifically, on a third of the trials, reward cues are presented simultaneously with advance task cues (i.e., by presenting the task cue in green font), and indicate the opportunity to earn monetary bonuses if performance is accurate and fast (relative to baseline performance) on that trial. By only presenting reward cues on a subset of trials, the remaining subset of non-incentivized trials and target stimuli can be directly compared across the proactive and baseline conditions. A divergence from Bugg & Braver (2015) is that single-task conditions are not included as part of the battery (due to length constraints), which precludes direct calculation of mixing costs. Nevertheless, the key prediction is that enhanced *proactive control* will lead to global performance improvement (i.e., present on all trials) even on these non-incentivized trials. In terms of cognitive control-related neural activity, the prediction is of increased cue-related activity, which might also be accompanied by sustained activation (in maintaining the reward incentivized motivational context) across the task block.

The *reactive* condition utilizes a new manipulation which has not previously been examined in prior work. Specifically, the reactive condition includes punishment (rather than reward) incentives, again on the same one-third of trials that were incentivized in the proactive condition. However, in the reactive condition the incentive cue is presented at the time of the target stimulus, rather than with the task cue (i.e., by presenting the target in green font), which prevents the use of incentive motivation in a preparatory fashion. Participants are instructed that they will lose a component of their potential monetary bonus if they make an error on these incentivized trials. Critically, the incentivized trials occur preferentially with incongruent target stimuli. This manipulation is intended to associate punishment-related motivation with these high-conflict items, potentially leading to increased response monitoring and caution when incongruence is detected. As such, the key prediction is that enhanced reactive control should reduce the TRCE, even on the non-incentivized trials, when compared to baseline and proactive conditions. In terms of neural activity dynamics, reactive control effects should be observable as transient probe-triggered activation on incongruent (relative to congruent trials).

### Sternberg WM

The Sternberg item-recognition task has been one of the most popular experimental paradigms used to assess short-term / working memory for over 50 years (Sternberg, 1966), but more recently has been adapted particularly for the study of cognitive control and in neuroimaging paradigms, with the “recent probes” variant (Jonides et al., 1998; Jonides & Nee, 2006). Like standard versions of the paradigm, the recent probes variant presents participants with a memory set of various load levels (number of items), followed after a short delay (retention period) by a single item probe, which requires a target response if the probe was a part of the memory set. However, in the recent probes variant, the key manipulation is that the probe item can also be a part of the memory set of the previous trial, but not the current trial, which is termed a “recent negative” (RN) probe. On these RN trials, the probe is associated with high familiarity, which can increase response interference and errors, unless cognitive control is utilized to successfully determine that the familiarity is a misleading cue regarding probe status (target or nontarget). Thus, a key index of cognitive control in this Sternberg variant is the *negative recency effect* (Jonides & Nee, 2006; Monsell, 1978), which contrasts RN and NN trials (NN: novel negative, when the probe item is not a member of the current or previous trial’s memory set; see Figure 1D).

The classic finding in the literature is that as the memory set increases in size (i.e., WM load increases) performance declines accordingly (Sternberg, 1969). Under conditions in which the WM load is below capacity (3-4 items), *proactive control* strategies can be utilized to keep the memory set accessible within WM, and used as an attentional template from which to prospectively match against the probe. In contrast, when the WM load is above capacity (∼7 items), *reactive control* strategies are likely to be utilized, with probe responses driven by retrieval-focused processes, such as monitoring of familiarity signals. The variants of the Sternberg WM included in the DMCC battery are adapted from prior studies, that utilized manipulations of both WM load expectancy (Speer et al., 2003) and RN frequency (Burgess & Braver, 2010). Specifically, in all conditions, trials randomly vary in set size, with words used as stimuli, such that all items are novel on each trial, with the exception of RN probes. Under such conditions, Burgess & Braver (2010) found strong RN interference effects are observed. Likewise, following Speer et al (2003), the set size in a given trial is revealed sequentially, leading to unpredictability and reliance on WM load expectancies to engage control strategies.

In the *baseline* condition, most trials have high WM load (6-8 items) and RN frequency is low, which should reduce tendencies to engage either proactive or reactive control strategies. However, in the *proactive* condition, most trials have low WM load (2-4 items), leading to the expectancy that active maintenance-focused and proactive attentional strategies will be effective, while RN frequency remains low (i.e., matched with the baseline condition), such that the utility of reactive control should be unchanged. The critical prediction concerns the 5-item set size which occurs equivalently in all conditions, and thus can be compared between them. The key hypothesis is that use of proactive control strategies, will improve performance, primarily for the target probe trials (termed novel positive, or NP, since they never overlap across trials). In terms of neural activity dynamics, the key prediction is of increased encoding-related activity (i.e., during presentation of the memory set) on all trials, which might also be accompanied by sustained activation (i.e., maintaining the attentional strategy) across the task block.

In the *reactive* condition, WM loads are identical to the baseline condition, while the frequency of RN trials is strongly increased. Thus, in the reactive condition, it is familiarity-based interference expectancy that increases, rather than WM load expectancy. Based on the increased interference-expectancy, the theoretical hypothesis is that participants will not rely on familiarity as a cue for responding, but will instead use familiarity cues to prompt evaluation of the probe match to items stored in WM. Consequently, the key prediction is that of reduced RN-related interference in the reactive condition. In terms of neural activity dynamics, reactive control effects should be observable as transient probe-triggered activation on RN (relative to NN trials).

## Methodological Approach

As this is the first paper to describe the DMCC project, we provide here a broad overview of the methodological approach. Our goal is to inform interested readers of the key project components and rationale, with the aim of promoting open science and reproducibility, and encouraging utilization of DMCC data within the scientific community. In the subsequent section, we focus on the methodological details specific to the first-set of analyses performed with this dataset.

### Participant Sample and Acquisition Protocol

In a number of ways, the DMCC project was modeled after HCP, and endeavored to follow closely to the tenets of the ‘HCP-style paradigm’ in terms of the participant sample and acquisition protocol (Glasser et al., 2016). In particular, like the HCP, the DMCC project is focused on examining both normative aspects and inter-individual variability in cognitive and brain function among healthy young adults. The sample was thus constrained to participants aged 18-45, which is an age range that is not strongly impacted by neurodevelopmental changes or by neurodegenerative changes associated with increasing age.

The scope and structure of the DMCC project imposed several important constraints, that caused us to deviate from the HCP paradigm in a number of ways that we describe here. To meet the demands of our project in isolating distinct components of cognitive control with high experimental precision, we required 25-30 minutes of scanning time for each of the 4 tasks. Since each of the three task fMRI sessions was structured to contain all 4 tasks, this resulted in ∼2 hours of task fMRI per session. Because of these constraints and the intense cognitive demands associated with performing our task battery, subject fatigue and motivation were found to be a significant challenge. As a solution to this challenge, we found it most helpful for fatigue and motivation recovery to schedule only a single imaging session per day and to space sessions out over a minimum of two days. This protocol of 3 spaced imaging sessions introduced strong logistical constraints that necessitated recruitment of participants from the HCP pool who were residents within the local community. This differentiated the DMCC from the HCP, which used a “burst scheduling” protocol to acquire data over a consecutive 2-day period, and enabled the recruitment of out-of-town family members.

Additionally, although it was a high priority to recruit family members into the DMCC, given the smaller scope of the DMCC project, and the challenges described in recruiting fully local twin pairs, our recruitment focus was on identical (monozygotic; MZ) twin pairs (and these pairs were primarily recruited from other sources than the HCP pool). Consequently, in the DMCC sample, there are very few DZ and non-twin siblings, which precludes true classical genetic modeling, which requires contrasts between MZ and DZ (or non-twin siblings) to separate genetic heritability from environmental effects. Nevertheless, the inclusion of MZ twin pairs are still highly valuable from a statistical modeling perspective, as they provide convergent information regarding both test-retest reliability (using twin similarity as a lower-bound estimate) of individual DMCC cognitive control measures as well as the ability to estimate the covariance structure among DMCC tasks that is not contaminated by individual-specific measurement error effects. As described briefly below, we have begun to fully exploit these properties using multivariate pattern analysis (MVPA) approaches.

After the start of data collection, the DMCC project was extended to incorporate an additional study design arm, supported by funding from the NCCIH. In this arm, MZ twins were recruited, at the time of initial screening, to participate in a mindfulness skills training intervention component. Following completion of their initial wave of testing, the twin-pair was randomly assigned, such that one twin received mindfulness training through an established and scientifically-validated 8-week course (Mindfulness-Based Stress Reduction), while the other twin served as a wait-list control. Following course completion, both twins returned for another wave of testing with the full DMCC protocol. The use of this novel, randomized discordant experimental twin design was another advantageous feature leveraging the utilization of MZ twins in the project.

The DMCC acquisition strategy was designed with the goal of facilitating integration of DMCC data with HCP data collected on the same participants. Thus, similar to the HCP, we utilized a multi-band acceleration sequence to increase spatial and temporal resolution in BOLD fMRI scans (Uğurbil et al., 2013). However, in line with recommendations from methodological evaluations (Todd et al., 2017), we adopted a more conservative approach similar to that of other contemporaneous projects (e.g., MyConnectome; Poldrack et al., 2015), by utilizing a multi-band acceleration factor of 4 (MB4), as opposed to the more aggressive 8-fold acceleration used in the HCP. This reduction led to a corresponding shift in acquisition parameters (2.4 mm^3^ voxel size, TR=1200 msec), but retained some of the gains in spatial and temporal resolution, while also maintaining high levels of signal quality with whole-brain coverage.

Participants in the DMCC protocol undergo a series of 4 experimental sessions. The first session involves out-of-scanner behavioral assessment geared towards providing a more comprehensive profile of individual difference characteristics. The assessment includes both self-report measures related to personality and psychological health and well-being, as well as cognitive tests of crystallized and fluid intelligence, processing speed, working memory capacity, and attentional control. A full-list and detailed description of these measures is beyond the scope of the current report, but can be found at this link (https://nda.nih.gov/edit_collection.html?id=2970). Here, we just briefly list a few of the notable measures and constructs being assessed: NIH toolbox (Oral Reading Recognition, Pattern Comparison, Flankers Task; Weintraub et al., 2013), Operation and Symmetry Span (Redick et al., 2012); Ravens Matrices (Raven, 2000); Five Factor Personality (NEO-FFI; McCrae & Jr, 2007); Trait Mindfulness (FFMQ, MAAS; (Baer et al., 2006; Brown & Ryan, 2003), Reward Sensitivity (BIS/BAS, SPSRQ; Carver & White, 1994; Torrubia et al., 2001), Emotion Regulation (ERQ; Gross & John, 2003), Need for Cognition (Cacioppo et al., 1984), Positive and Negative Affect (PANAS; Watson et al., 1988), State and Trait Anxiety (STAI; Spielberger, 2010), Sleep Quality (Pittsburgh Sleep Quality; Buysse et al., 1989), Impulsiveness (Patton et al., 1995), and Life Satisfaction (SWL; Diener et al., 1985).

Following the behavioral session, participants always completed the Baseline neuroimaging session first, then the Proactive and Reactive neuroimaging sessions, with the order of these latter two sessions counter-balanced across participants. Within the session, task-order was counter-balanced but maintained across all subsequent sessions, and testing waves, for longitudinal participants. Likewise, in twin-pairs, both members of the pair experienced the same task and session order, in order to facilitate twin-based comparisons. In addition to these behavioral and neuroimaging data, relevant physiological information is also collected during the sessions, including weight, blood pressure, heart rate, caffeine and food intake, plus a number of salivary cortisol samples (including early morning, pre and post scanning) and collection of DNA on all participants. During scanning, heart-rate and respiration data are also continuously monitored and collected, with the potential for use in preprocessing.

### Analysis Workflow

Given the complexity of the DMCC data acquisition protocol, and the amount of data generated per participant, we found it critical to develop a set of standardized operating procedures and processing pipelines, that enable efficient and high-levels of quality control of a project of this type, while also allowing for a de-centralized means of management, familiarization, and processing by multiple members of the research team (Etzel et al., 2020). Moreover, each component of the processing pathway and associated software tools were developed with the aim of maximizing the transparency, reproducibility, and the data sharing potential of the project. Data storage and management are structured to enable both secure archiving as well as tight integration with associated data from the HCP, by making use of the IntraDB databasing file format (Marcus et al., 2013), and the Connectome Coordination Facility for data structures and storage capabilities (https://www.humanconnectome.org/study/dual-mechanisms-cognitive-control). Behavioral (and later neuroimaging) data are also being deposited in the NIH National Data Archive and Repository (NDAR), with both repositories (NDAR and CCF) being used to facilitate data sharing. Currently, a small sample of the DMCC neuroimaging dataset is available on OpenNeuro at the following link (https://openneuro.org/datasets/ds002152/versions/1.0.2). Schematic diagrams of DMCC workflows along with associated documentation on standardized operating procedures, processing scripts, software tools, and example outputs are all available on OSF, and can be accessed via the following link (https://osf.io/xfe32/).

Because of our primary interest in individual differences in the DMCC project, we developed an analysis workflow that could provide a semi-automated and reproducible processing pipeline that generates quality control information as well as easily visualized summaries of all processing stages at the single-subject level. For our initial set of analyses, we adopted the HCP pipeline approach to preprocessing (Glasser et al., 2013), which takes advantage of the high spatial resolution and multi-modal imaging data to enable artifact removal, distortion correction, and align brain areas. Critically, we also examined that potential benefits of surface-based registration and reconstruction, using CIFTI file-formats developed for the HCP.

A key aspect of our approach has been to leverage cortical parcellation atlases as a primary analytic tool. In particular, our approach to single-subject analysis, utilizes the cortical parcels (and subcortical nuclei) as the basic unit of analysis. For univariate analyses, this is done by averaging voxel-wise data within parcels (nuclei), which results in a highly significant reduction of data complexity, and enabling an effective way to visualize whole-brain data in a principled yet unbiased manner. For multivariate analyses, the voxel-level data are retained, but analyses are conducted at the parcel (nuclei) level, treating these as an unbiased / pre-specified means of conducting whole-brain or region-of-interest analyses. We have explored a number of these cortical parcellation atlases, including Multi-Modal Parcellation (MMP; Glasser, Coalson, et al., 2016), but for our primary set of analyses, we utilized Schaefer 400-parcel atlas, as it uses a combined global-local approach (gradient-weighted Markov Random Field) to create more homogenously sized parcels, available at various resolutions (Schaefer et al., 2018). Additionally, we include a set of 19 segmented subcortical nuclei (e.g., amygdala, hippocampus, caudate, etc), generated as part of the CIFTI file format in the HCP pipeline (Glasser et al, 2013).

Throughout the project we have been committed to conducting methodological evaluations and benchmarking to ensure that we are following best practices and incorporating state-of-the-art approaches throughout. As such, we found that surface-based alignment, combined with the cortical parcellation approach, exhibited significant benefits in statistical sensitivity to task effects relative to more standard volume-based alignment approaches; consequently, the data and analyses reported here utilize this approach. Additionally, we are now in the process of shifting our pre-processing pipeline to fMRIprep (Esteban et al., 2018, 2020), as we have that it yields further benefits in terms of standardization, QA/QC, and statistical sensitivity, relative to the HCP pipelines (Etzel et al., 2019). Most recently, we have explored a second stage pre-processing approach using a whole-brain neural model derived from resting-state fMRI scans to filter out intrinsic activity dynamics and provide even greater sensitivity and temporal precision regarding event-related activity dynamics (Wang et al., 2020). Finally, we have been focusing on multivariate pattern analysis (MVPA) approaches (both classification-based decoding and representational similarity analyses; RSA) as a more powerful and flexible means from which to fully leverage the advantages of our twin-based design (Etzel et al., 2020), to identify task-related activity, and to reveal key neural coding properties (Freund et al., 2020; Freund & Braver, 2020; Tang et al., 2020) All of these benchmarking and comparative analyses will be the focus of forthcoming reports; here, we report standard univariate analyses based on Schaefer atlas cortical parcellations using the HCP surface-based processing pipelines.

For univariate analyses, standard GLM estimation is conducted (using AFNI software) according to a mixed blocked / event-related design (Petersen & Dubis, 2012; Visscher et al., 2003). This design and the associated analytic approach are important for testing the DMC framework, as this approach allows for the separation of sustained (block-related) from transient event-related activation dynamics. Many theoretical predictions regarding dissociations between reactive and proactive control involve distinctions between transient probe-related activity (reactive effects) from either cue-related or sustained activations (proactive effects). The mixed blocked / event-related GLM analyses are facilitated by the consistent structure present in task fMRI scanning runs, which each comprise three task blocks (of approximately 3-minute duration) alternating with four resting fixation blocks (30-seconds duration each). Likewise, within each task block, inter-trial intervals are jittered in a uniform manner to improve estimation of event-related effects. Finally, event-related activation is estimated using a deconvolution (finite impulse response, FIR; Glover, 1999) approach. This approach enables flexible estimation of the full timecourse of event-related activity which is useful in multi-event trials (Ollinger et al., 2001), such as the cue-target structure present in 3 of the 4 DMCC tasks (AX-CPT, Cued-TS, Sternberg).

Following GLM estimation, data are summarized for visualization at parcel/nuclei level, in terms of sustained estimates and event-related timecourses for each condition and task and condition, breaking the event-related data down further into different trial types and contrasts. From these, summary reports are generated for each participant (which are also linked to parallel summaries of the in-scanner behavioral performance data), enabling easy visual inspection and quality control checks. An illustrative example of these single-subject summary reports can be found at the following link (https://osf.io/54qhb/). A further strength of this analytic and visualization approach is that it facilitates exploration and benchmarking of the effects of various processing parameters, parcellation atlases, and pipeline choices as described above.

## Focus of the Current Report

Because of the rich nature of the DMCC dataset and the on-going state of data collection, many primary research questions of interest will be the focus of future papers. Most prominently, individual difference focused analyses, which will examine the relationship between neural activity patterns associated with proactive and reactive control, and the other individual difference measures, will be most powerfully investigated after data collection is complete (estimated 2023), with a target sample size of 200. Here we focus our initial analyses on the question of whether there is a unique neural signature of reactive control, that is distinct from proactive control. Although proactive-control focused analyses are equally important, the examination of reactive control has been somewhat neglected in ours and other researchers’ prior work. Consequently, we took advantage of the within-subject design and novel DMCC task battery, with 4 distinct tasks including baseline and reactive control conditions to test whether there are consistent reactive control patterns across the task, and whether these are linked to behavioral performance enhancements associated with increased utilization of reactive control.

## Methods

### Participants

Analyses were conducted on a subset of 80 participants (Mean Age=32.0, SD=6.1, Female=47, Male=32, Prefer Not To Answer=1) with high-quality data from both baseline and reactive sessions. This participant sample included both prior HCP participants (N=43) as well as 24 twin-pairs, and had the following self-reported demographic breakdown: White/Caucasian (57), Black/African-American = 11, Asian / Pacific Islander = 5, More Than One Race = 5, Prefer Not To Answer = 1. During recruitment all participants were screened and excluded if they were outside the 18-45 age range, were diagnosed with severe mental illness, significant neurological trauma, certain medication usage, or failure to pass screening for MRI safety contraindications. Participants recruited from the HCP sample were screened and excluded for a significant change in their status with regard to these exclusion criteria from the time of their last HCP scan. A screening form with the full listing of exclusion criteria can be found at the following link (https://osf.io/6efv8/). All participants provided written informed consent in accordance with the Institutional Review Board at Washington University, St. Louis, and received $400 compensation for completion of all sessions (along with additional monetary bonuses for performance in the Cued-TS sessions).

### Imaging Data Acquisition

All imaging data were acquired on a Siemens 3T PRISMA scanner using a 32-channel head coil, and included both high-resolution MPRAGE anatomical scans (T1- and T2-weighted with 0.8 mm^3^ voxels) and BOLD functional scans using the CMRR multi-band sequences described above (acceleration factor=4, 2.4 mm^3^ voxels, 1.2s TR, with alternating AP and PA encoding directions). Full details of the acquisition protocol and parameters can be found at the following link (https://osf.io/tbhfg/). In each of three imaging sessions, participants underwent 8 BOLD scans runs of approximately 12 minutes in duration, during which they performed 2 consecutive runs of each of the 4 DMCC tasks.

#### Tasks

All DMCC tasks were programmed and presented to participants using Eprime software (Version 2.0, Psychology Software Tools, Pittsburgh, PA). All tasks scripts are available at the following link (http://pages.wustl.edu/dualmechanisms/tasks). In all tasks but the Stroop, participants responded with a custom designed manual response box, using index and middle fingers of the right hand. In the Stroop, participants made vocal responses that were digitally recorded and later processed to automatically extract response latencies, using a MATLAB spectral filtering algorithm, with code available at the following link (https://github.com/ccplabwustl/dualmechanisms/tree/master/preparationsAndConversions/audio). These recordings were also manually inspected for quality control purposes, and to code response errors.

In the *Stroop* task (see Figure 1A), participants named the font color of the word (red, purple, blue, yellow, white, pink, black or green). The key contrast of interest is of biased incongruent vs. biased congruent trials. In the *AX-CPT* task (Figure 1B), participants made target or non-target button press responses to cue and target stimuli, with target trials defined as an A cue followed by an X probe. The key contrast of interest is BX vs. BY trials. In the *Cued-TS* paradigm (see Figure 1C), participants made target or non-target button press responses to letter-digit target stimuli, with the target defined by the task cue (Letter or Digit) and response mapping (Letter: consonant or vowel; Digit: odd or even). The key contrast of interest is non-incentive incongruent vs. non-incentive congruent trials. In the *Sternberg* paradigm (see Figure 1D), participants made target or non-target button press responses to probe word stimuli, deciding whether they had been one of the words previously presented in the memory set for that trial. The key contrast of interest is recent negative critical (5-item) trials vs. novel negative critical (5-item) trials. Detailed information regarding stimulus parameters is available at the following link (https://osf.io/48aet/).

#### Data Analysis

All analyses were performed using R statistical software version 3.1.3 (R Development Core Team 2015). Code and input data for replicating the figures and statistics are available at the following link (https://osf.io/xvzrf/).

For behavioral data, analyses focused on RT and error rates in the key contrasts of interest. These contrasts isolate interference effects associated with increased cognitive control demands, or in other words, the degree to which task performance declines in high control relative to low control trials. Second-level (i.e., group effect) statistical analyses were conducted using robust t-tests (Yuen), which operate on trimmed means (0.1 trim) to reduce the effects of outliers (Yuen, 1974).

For imaging data, preprocessing was implemented with HCP preprocessing pipelines (Glasser et al., 2013), which utilize FreeSurfer to reconstruct the cortical surface and extract subcortical nuclei (Fischl et al., 2002), align the data to MNI atlases in surface-space, correct distortions, compute motion alignment parameters, introduce spatial smoothing, and express the data in “grayordinate” vertices. Following HCP preprocessing, additional pre-processing occurred with AFNI software (Cox, 1996), computing frame-wise motion censoring parameters (FD > 0.9 mm) and implementing image normalization (i.e., demeaning). Sustained and event-related data were estimated using standard AFNI GLM procedures (3dREMLfit) at the voxelwise level (which includes 6-parameter motion and framewise censoring parameters, as well as polynomial detrending, using the -polort flag). Following GLM estimation, voxelwise beta estimates were averaged into cortical parcels according to the Schaefer 400 atlas (Schaefer et al., 2018), and subcortical parcels via the CIFTI FreeSurfer segmentation (19 nuclei) (Glasser et al., 2013). For the primary analyses of interest, contrast timecourses were computed, and extracted around the 2-TR peak of the target event (allowing for the ∼5 second hemodynamic lag). These contrasts isolate interference effects associated with increased cognitive control demands, or in other words, the degree to which neural activity increases on high control relative to low control trials. Second-level (i.e., group effect) statistical analyses were performed to test whether a contrast was significant (i.e., greater than zero, indicating high control demand activity > lower control demand activity) using 1-sample t-tests.

Participants were included in analyses if they successfully completed all experimental sessions. Of the 80 participants meeting these criteria for the initial analyses, 10 participants did not have full imaging data for all Baseline and Reactive scans (most were missing one scanning run due to technical issues). Additionally, 14 participants were missing behavioral data during at least one scanning run (again due to technical issues, typically related to button box or microphone malfunction). Because of the loss in statistical power, GLM estimation and behavioral data analyses were not conducted for these participants, for the problematic task in the relevant session.

## Results

### Baseline Session: Behavioral Performance

We examined the presence of interference effects in the Baseline session, associated with high vs. low cognitive control demands, through the relevant contrast of interest for each task, in both reaction time and error rates. In all four tasks, and in both behavioral measures, these interference effects were highly robust (see Table 1): Stroop (incongruent–congruent; RT: t = 18.37, p < .001; error: t = 4.49, p < .001), AX-CPT (BX-BY; RT: t = 8.78, p < .001;error:t = 7.46, p < .001), Cued-TS (incongruent–congruent; RT: t =5.34, p <.001; error: t = 5.44, p < .001), Sternberg (RN–NN; RT: t = 7.30, p < .001; error: t = 9.94, p < .001). These results are consistent with the interpretation that the selected trial type contrasts index increased cognitive control demands.

**Table 1.** Behavioral measures for high and low control demand trials in DMCC tasks. Left columns indicate Baseline condition, right columns indicate Reactive condition. Data are reported as means with SEM in parentheses, for both reaction time (RT) and error rate.

|  | Baseline |  |  |  | Reactive |  |  |  |
| --- | --- | --- | --- | --- | --- | --- | --- | --- |
|  | High RT | Low RT | High Error | Low Error | High RT | Low RT | High Error | Low Error |
| Stroop | 930 (16) | 795 (16) | 0.024 (0.005) | 0.003 (0.001) | 855 (18) | 771 (16) | 0.019 (0.004) | 0.002 (0.002) |
| AX-CPT | 610 (25) | 466 (14) | 0.169 (0.02) | 0.028 (0.006) | 584 (22) | 430 (13) | 0.103 (0.02) | 0.033 (0.006) |
| Cued-TS | 1161 (32) | 1108 (32) | 0.076 (0.01) | 0.028 (0.004) | 1079 (37) | 1016 (33) | 0.045 (0.01) | 0.02 (0.003) |
| Sternberg | 1109 (29) | 946 (22) | 0.266 (0.02) | 0.055 (0.01) | 1033 (25) | 909 (20) | 0.192 (0.02) | 0.041 (0.02) |

### Baseline Session: Neural Activity

We conducted a whole-brain parcel-wise analyses to identify regions showing robust effects of cognitive control demand. These analyses were based on event-related timecourses for the high – low cognitive control demand contrast in each task, isolating a 2-TR peak window around the target event (assuming a ∼5-second hemodynamic lag, and with the peak selected based on visualization of initial pilot data; see Figure 4). A two-stage conjunction analysis approach was then utilized to reveal regions showing consistent control demands across all four tasks. In the first-stage, a linear mixed effect statistical model was employed to test each parcel for the fixed effect of high > low control demand, including the 4 tasks and participant as random effects. For this first-stage, to control for multiple comparisons a Bonferroni correction was used to control for the number of parcels (419). Thus, the significance level was set at p < .0001. In the second stage, each task was interrogated individually, for the significance of the contrast, using a p < .05 uncorrected significance levels. We counted a parcel as robust and consistently activated by control-demand if it not only passed the first stage, but was also significant for each of the 4 tasks, in the second stage.

From these analyses, we identified a set of 34 cortical parcels (no subcortical nuclei were identified) that met both criteria. These parcels are shown in Figure 2, with contrast timecourses shown in Figure 4. The identified parcels are primarily localized to regions within the frontoparietal and cingulo-opercular networks typically associated with cognitive control (Cole & Schneider, 2007; Dosenbach et al., 2008; Vincent et al., 2008). Indeed, 28 of the 34 regions are located in parcels labeled in the Schaefer atlas as belonging to either the Control (12), Salience/VentralAttention (11), or DorsalAttention networks (note that the 4 parcels labeled as belonging to the Default Mode network were located within lateral prefrontal cortex; see Table 2). The most robustly activated regions were bilateral, within dorsolateral prefrontal cortex (parcel #’s 140,347) and anterior insula / frontal operculum (parcel #’s 99, 306). These results imply that this set of regions may function as core network consistently responsive to cognitive control demands, across a wider set of task conditions.

**Figure 2.**
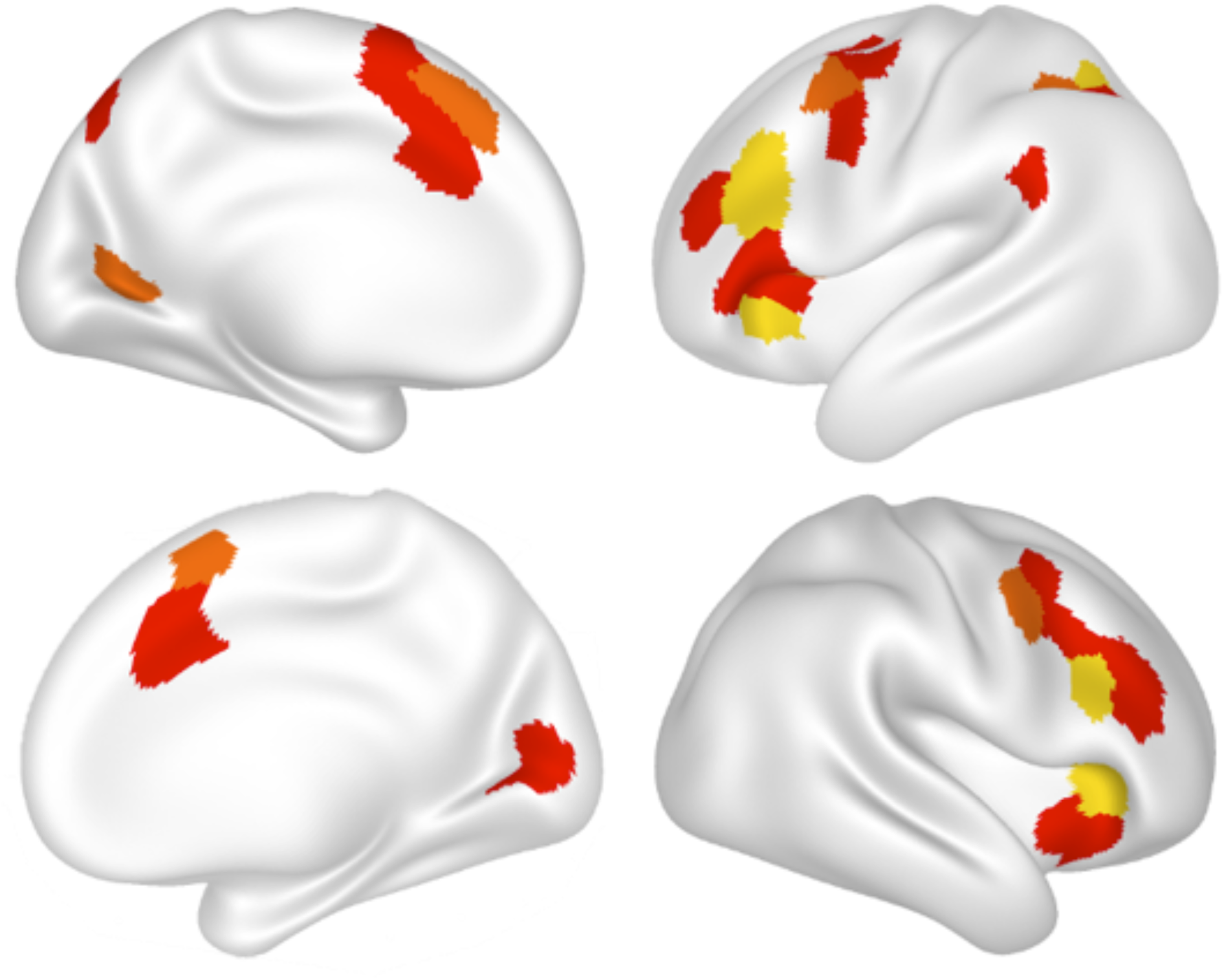
Brain regions identified showing a consistent effect of cognitive control demand in each of the four DMCC tasks. Regions are shown as surface-based parcels in the Schaefer 400 atlas (Schaefer et al, 2018) shown on medial (left side) and lateral (right side) surfaces, for the left (upper) and right (lower) hemispheres. Color scale indicates contrast significance in each of the 4 tasks: red (min t > 1.96; p <∼ .05), orange (min t > 2.57; p < ∼ .01), and yellow (min t > 3.0; p < ∼ .005). Regions are mostly localized to fronto-parietal and cingulo-opercular network (see Table 2 for anatomical locations).

**Table 2.** The 34 parcels identified as showing a consistent effect of cognitive control demand across DMCC tasks. Parcel numbers, names, networks, and centroid coordinates (MNI) are those provided in the 400 parcel, 7 network resolution Schaefer atlas (Schaefer et al, 2018). For additional information, see https://github.com/ThomasYeoLab/CBIG/tree/master/stable_projects/brain_parcellation/Schaefer2018_LocalGlobal.

| Parcel Number | Hemisphere | Network | ROI | X | Y | Z |
| --- | --- | --- | --- | --- | --- | --- |
| 22 | Left | Visual | 22 | -19 | -65 | 7 |
| 77 | Left | Dorsal Attention | Post_9 | -33 | -46 | 41 |
| 78 | Left | Dorsal Attention | Post_10 | -29 | -58 | 50 |
| 86 | Left | Dorsal Attention | FEF_1 | -40 | -3 | 51 |
| 87 | Left | Dorsal Attention | FEF_2 | -25 | -1 | 55 |
| 91 | Left | Dorsal Attention | PrCv_2 | -50 | 3 | 38 |
| 93 | Left | Salience/Ventral Attention | ParOper_2 | -58 | -44 | 27 |
| 99 | Left | Salience/Ventral Attention | FrOperIns_3 | -33 | 25 | -1 |
| 101 | Left | Salience/Ventral Attention | FrOperIns_5 | -33 | 19 | 8 |
| 103 | Left | Salience/Ventral Attention | FrOperIns_7 | -43 | 12 | 2 |
| 105 | Left | Salience/Ventral Attention | FrOperIns_9 | -52 | 9 | 13 |
| 107 | Left | Salience/Ventral Attention | Med_1 | -6 | 22 | 31 |
| 110 | Left | Salience/Ventral Attention | Med_4 | -5 | 9 | 48 |
| 127 | Left | Control | Par_1 | -29 | -74 | 42 |
| 130 | Left | Control | Par_4 | -35 | -62 | 48 |
| 139 | Left | Control | PFCI_5 | -42 | 38 | 22 |
| 140 | Left | Control | PFCI_6 | -45 | 20 | 27 |
| 144 | Left | Control | pCun_1 | -9 | -77 | 45 |
| 148 | Left | Control | PFCmp_1 | -4 | 28 | 47 |
| 172 | Left | Default | PFC_7 | -48 | 28 | 0 |
| 175 | Left | Default | PFC_10 | -53 | 19 | 11 |
| 185 | Left | Default | PFC_20 | -42 | 7 | 48 |
| 189 | Left | Default | PFC_24 | -6 | 10 | 65 |
| 219 | Right | Visual | 19 | 9 | -74 | 9 |
| 301 | Right | Salience/Ventral Attention | PrC_1 | 51 | 3 | 41 |
| 303 | Right | Salience/Ventral Attention | FrOperIns_2 | 41 | 8 | -3 |
| 306 | Right | Salience/Ventral Attention | FrOperIns_5 | 37 | 23 | 5 |
| 314 | Right | Salience/Ventral Attention | Med_4 | 6 | 11 | 58 |
| 340 | Right | Control | PFCv_1 | 34 | 21 | -8 |
| 346 | Right | Control | PFCI_6 | 50 | 30 | 18 |
| 347 | Right | Control | PFCI_7 | 48 | 18 | 23 |
| 349 | Right | Control | PFCI_9 | 47 | 29 | 28 |
| 350 | Right | Control | PFCI_10 | 39 | 11 | 34 |
| 353 | Right | Control | PFCI_13 | 43 | 7 | 51 |

### Baseline: Brain-Behavior Relationships

Given that both behavioral performance and parallel neural activity measures showed consistent effects of control demand across the four tasks, we next tested whether these effects were associated. In particular, we hypothesized that individuals showing larger behavioral interference effects would also show larger neural activity effects of control demand, i.e., a positive correlation. To test this hypothesis, and reduce dimensionality, we created a composite behavioral index of performance, by z-score normalizing the data across participants, and then combining RT and error measures across the 4 tasks to create a summed z-score value for each participant, with larger values indicating higher interference. Likewise, to create a single neural activity index, we treated the 34-parcels as a “mega-parcel”, averaging contrast activity values (betas) across the set. In addition, we z-score normalized, and then summed these mega-parcel values across the four tasks for each participant. We then correlated the neural activity index against the behavioral composite index. This correlation was significantly positive (r: .25; Pearson’s: p = .04; Spearman’s: p = .03; see Figure 5A), supporting our hypothesis.

### Reactive Effects: Behavioral Performance

We next examined whether the experimental manipulations designed to encourage utilization of reactive control were effective in enhancing behavioral performance by reducing interference effects. Paired robust (Yuen) t-tests were conducted to compare RT and errors across Baseline and Reactive sessions, for each of the tasks. In 3 of the 4 tasks, at least one of the behavioral metrics showed a statistically significant effect, and this was trending in the remaining task: Stroop (RT: t = 8.01, p < .001; error: t = 1.32, p > .1), AX-CPT (RT: t = -1.23, p > .1; error: t = 3.22, p = .002), Cued-TS (RT: t = 0.1, p > .1; error: t = 1.88, p = .06), Sternberg (RT: t = 1.67, p = .1; error: t = 2.32, p = .02). Thus, behavioral data support the hypothesis of enhanced task performance (reduced interference) via increased utilization of reactive control (see Figure 3). To support this interpretation, we conducted a paired robust (Yuen) t-test on the behavioral composite index aggregated across the 4 tasks, which was computed separately for Reactive as well as Baseline, for each participant. This analysis confirmed the hypothesis of a robust decrease in interference (improved cognitive control utilization) in Reactive, relative to Baseline (t = 5.36, p < .001).

**Figure 3.**
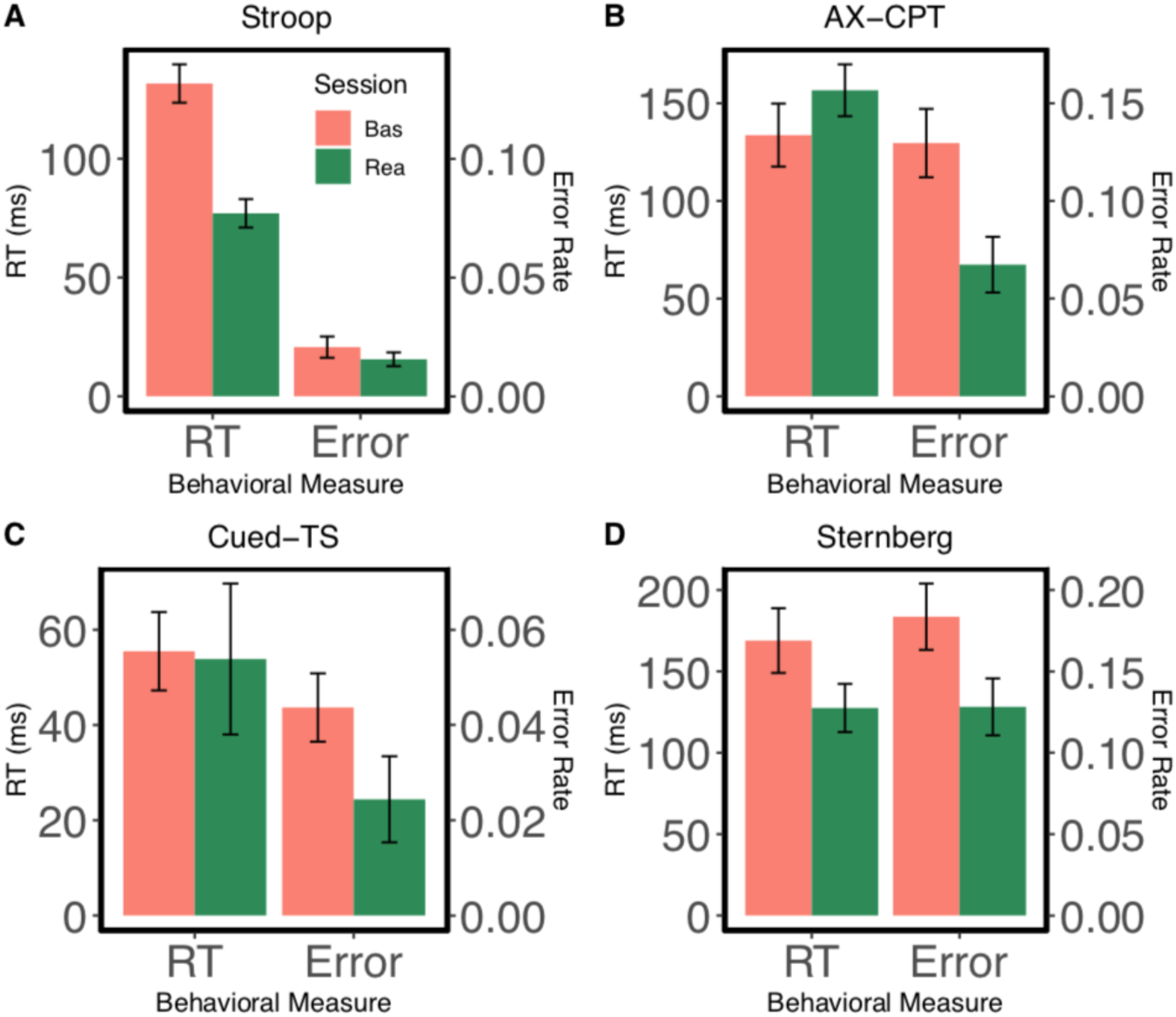
Effects of Reactive condition on behavioral performance in DMCC cognitive control indices. Data are shown in terms of RT (left side / axis) and error rate (right side / axis) for each of the four DMCC tasks: A. Stroop; B. AX-CPT; C. Cued-TS; D. Sternberg. Baseline (red bars) and Reactive (green bars) data shown separately. Reductions in cognitive control indices indicate decreased interference in each task, consistent with enhanced control in the Reactive condition.

### Reactive Effects: Neural Activity

We examined whether performing the four tasks under Reactive control conditions would be associated with parallel neural activity effects to what was observed in terms of behavioral performance. Visualization of the event-related contrast timecourses for each task suggested a reduction in activation in Reactive, relative to Baseline (see Figure 4) in the 34-megaparcel. Paired robust (Yuen) t-tests were conducted to statistically compare neural activity during the target period across Baseline and Reactive sessions, for each task, in the 34-megaparcel. In all four tasks, we found that activity was numerically reduced in the Reactive session, and this reduction effect was statistically significant in 3 of the 4 tasks: Stroop (t: 3.83, p < .001), AX-CPT (t: 0.53, p > .1), Cued-TS (t: 2.16, p = .02), Sternberg (t: 2.93, p = .005). Thus, the neural data support the interpretation of reduced activation associated with increased utilization of reactive control. To support this interpretation, we conducted a paired robust (Yuen) t-test on the mega-parcel, by again computing a neural activity index averaging across the four tasks. This analysis strongly supported the pattern of a robust decrease in activation in Reactive, relative to Baseline (t = 4.58, p < .001).

**Figure 4.**
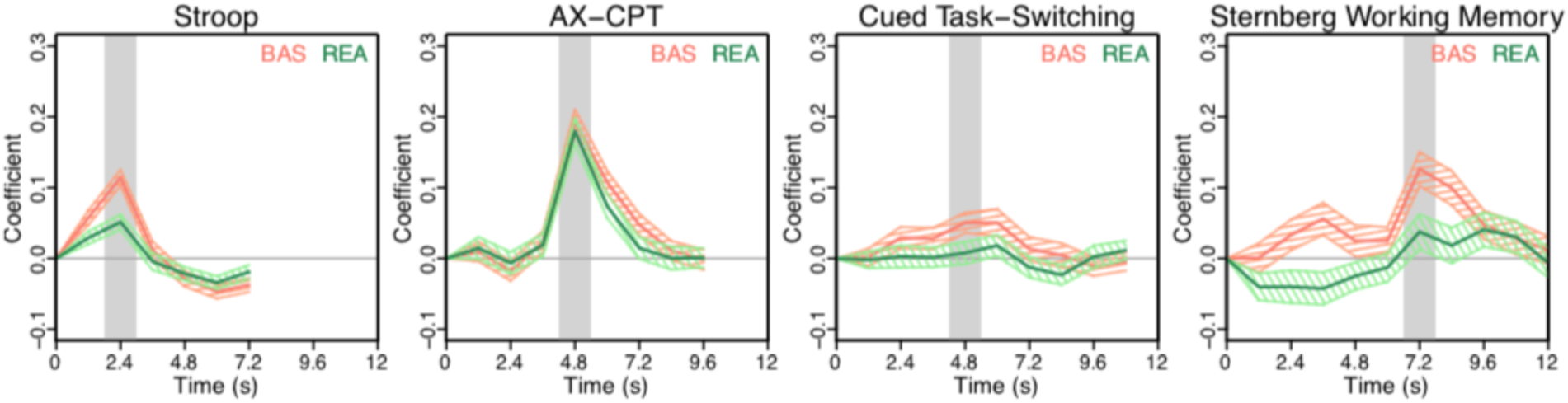
Event-related contrast effects for the DMCC tasks in Baseline and Reactive conditions. Data are shown for the 34-megaparcel as contrast timecourses (i.e., high – low control demand differences), with the key target period (2 TR window) shaded in gray. In the Baseline condition (red lines) there is a clear peak for each of the four DMCC tasks. This peak is consistently reduced across tasks in the Reactive condition (green lines).

### Reactive Effects: Brain-Behavior Relationships

We hypothesized that the Reactive-related enhancement of task performance (reduced interference effects) might be functionally linked to the parallel reduction of neural activity observed in fronto-parietal regions (34-megaparcel). To test this hypothesis, we first computed the brain-behavior correlation within the Reactive condition by itself, using the behavioral composite index and neural activity index. Interestingly, within the Reactive condition, we did not observe a significant brain – behavior relationship (r = .10; Pearson’s p > .1, Spearman’s p > .1; see Figure 5B). We next computed Baseline – Reactive change scores on these indices. Here, the brain-behavior correlation between these change scores was significant (r = .32; Pearson’s p = .01, Spearman’s p = .01; see Figure 5C). A more formal test of the hypothesis was conducted as a within-subjects statistical mediation analysis, according the procedures outlined in Judd et al. (2001). That is, we tested whether the effect of the Reactive experimental manipulation on behavioral performance was statistically mediated (or moderated) by its effect on reducing neural activity. This analysis did find evidence of statistically robust mediation (estimate: 0.42, t = 2.77, p = .007) without evidence of moderation (estimate: -.21, t: -1.53, p > .1).

**Figure 5.**
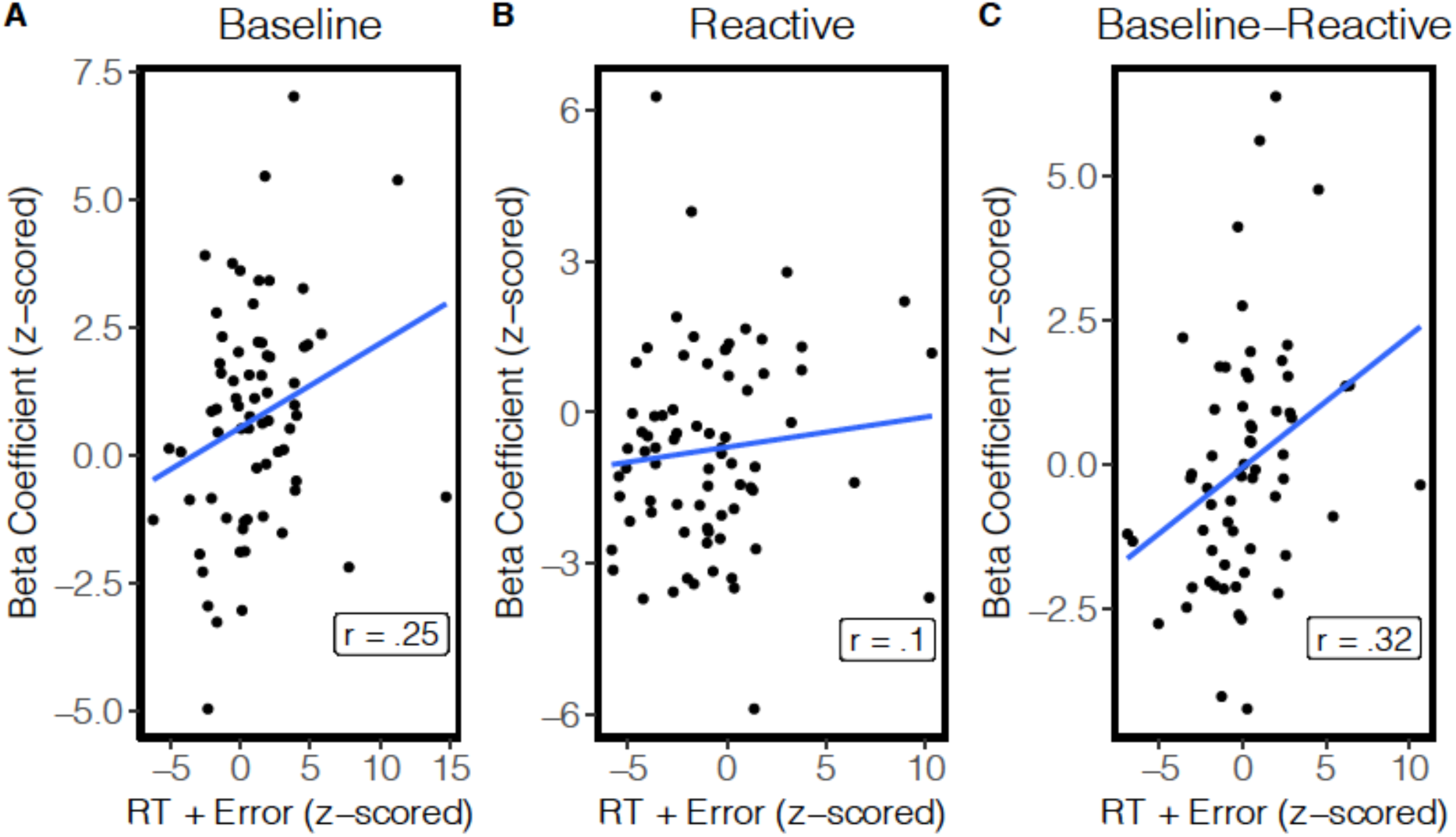
Brain-behavior correlations in Baseline and Reactive conditions. Scatterplot data show each participant in terms of their neural activity index (z-normalized beta coefficient for the 34-megaparcel summed across the 4 tasks) and behavioral composite index (high – low demand cognitive control indices for RT and error z-normalized and summed across the 4 tasks). The significant positive correlation in Baseline (A) indicates that individuals showing greater control-related activity also showed more behavioral interference. This relationship was not significant in the Reactive condition (B), but was significant in terms of the Baseline – Reactive change score (C). The positive correlation in the change score indicates that individuals showing a larger Reactive-related reduction in activity also tended to show a larger Reactive-related reduction in behavioral interference (i.e., enhanced control).

### Control Analyses

A confound present in the above analyses is that the Reactive experimental session always occurred after the Baseline session. This was an intentional aspect of the study design, to enable the Baseline session to provide an unbiased estimate of individual differences, and provide a practice foundation for the subsequent sessions. However, it does leave open the interpretation that the observed Reactive effects were a by-product of this systematic order relationship (e.g., practice or habituation related). Although it is impossible to fully resolve this potential confound, we did conduct two control analyses to address it. First, there was counterbalancing across participants, such that the Reactive session could be performed either second or third. We examined whether session order strengthened the Reactive effect, predicting greater effects when Reactive was the third session, if these were primarily due to order. However, we observed no difference in either the neural activity index (t: -1.27, p > .1) or the behavioral composite index (t: -1.4, p > .1). Likewise, we tested whether the Baseline – Reactive brain-behavior correlation was impacted by whether the Reactive condition was in the second or third session. There was no impact of this variable, and both correlations were of similar magnitude (reactive 2^nd^: correlation = .31; reactive 3^rd^: correlation = .35).

## Discussion

The focus of this report is twofold. First, we provide an overview of the DMCC project, with its approach to investigate cognitive control function in a theoretically-driven and experimentally-rigorous manner, with a focus on individual differences and brain-behavior relationships. Second, we highlight the promise and potential of this project for researchers interested in the neural mechanisms of cognitive control, by reporting initial analyses of DMCC data that reveal a novel neural signature of reactive control, in terms of decreased stimulus-triggered activity within a set of fronto-parietal brain regions. We elaborate on each of these dimensions in turn.

### DMCC Project Features

The primary feature of the DMCC study is its within-subjects design, in which each participant undergoes over 5 total hours of task fMRI, performing 4 distinct, but theoretically-targeted cognitive control paradigms (Stroop, AX-CPT, Cued-TS, Sternberg) under three different control modes (baseline, reactive, and proactive). There are now a number of other studies, utilizing much larger sample-sizes than the DMCC, in which each participant undergoes fMRI scanning while performing multiple cognitive tasks (e.g., HCP, ABCD, IMAGEN, PNC; Casey et al., 2018; Essen et al., 2013; Satterthwaite et al., 2014; Schumann et al., 2010) as well as smaller studies that have collected comparable task fMRI data on each individual (Gordon et al., 2017; Nakai & Nishimoto, 2020). However, to our knowledge, the DMCC project is the first collect this much task fMRI data within the domain of cognitive control, using theoretically-targeted experimental paradigms. Moreover, the use of a multi-task approach aligns well with psychometric and measurement model perspectives, in which the use of multiple task indicators assessing the same theoretical construct provides the foundation for more statistically valid extraction of latent variables (Conway & Kovacs, 2013; Cooper et al., 2019; Friedman & Miyake, 2016; Kievit et al., 2011). In other words, by collecting data on each participant from multiple cognitive control tasks, the DMCC project provides a more psychometrically valid and meaningful basis for determining whether neural and behavioral markers reliably tap into cognitive control, in showing a consistent pattern across tasks. Moreover, from a construct validation perspective, it can be argued that the DMCC data most strongly demonstrate the generalizability of behavioral and neural markers of cognitive control constructs, by minimizing the impact of measurement error or anomalous / spurious effects that might be observed within single tasks.

A second key aspect of the DMCC design is that not only do participants perform multiple cognitive control tasks under multiple experimental contexts, but that these contexts were also theoretically-designed to manipulate the mode of cognitive control being deployed. In particular, the DMC framework suggests a meaningful distinction between proactive and reactive control modes, in that these can be distinguished in terms of operating characteristics, temporal dynamics, and neural coding schemes. The DMCC project provides an unprecedented opportunity to systematically test this theoretical framework, by examining the neural activity and behavioral profiles of individuals performing each of the four tasks in the DMCC battery under proactive and reactive, as well as baseline conditions. Moreover, each of the individual tasks provides a rich set of data regarding the effects of different experimental manipulations on cognitive control modes and mechanisms, such as interference expectancy, instructed strategies, motivational incentives, and working memory load. Conversely, when examined in combination, the dataset enables a test of whether consistency in control mode shifts occur even across heterogenous tasks and experimental contexts. As such, the DMCC project can also be construed as a large test-retest design, from which state-dependent components of cognitive and neural activation profiles can be uncovered and disentangled from the state-independent processes that are engaged as individuals perform demanding cognitive tasks. As discussed further below, here we provide the first analysis of this type, leveraging the advantages of the DMCC to demonstrate the presence of consistent state-dependent changes in neural activity profiles occurring during reactive control.

A third key aspect of the DMCC design is its focus on individual differences, enabling examination of cognitive control variation in relation to other established sources of individual variation. Because of the broad range of individual difference measures collected outside of the scanner, it will be possible to relate variation in cognitive control function and control modes with a range of related constructs, including personality traits (e.g., anxiety, neuroticism, impulsivity, reward and punishment sensitivity, need for cognition), as well as with closely linked cognitive dimensions such as fluid intelligence, working memory capacity, and processing speed. Another source of individual differences information is the HCP, since a large subset of DMCC participants also took part in the HCP. Consequently, it is possible to leverage the set of task fMRI paradigms used in the HCP, as well as its larger sample-size (> 1000 participants), to draw inferences and connections regarding neural substrates. Moreover, the larger sample size and genotyping available in the HCP make it possible to link neural and behavioral cognitive control profiles identified in the DMCC with relevant dimensions of genetic variation. Additionally, the subset of MZ twins within the DMCC sample makes it possible to examine the effects of heritability and shared environmental on cognitive control variation in a manner that is more systematic and theoretically-driven than has been possible in prior investigations within this domain.

The longitudinal component of the DMCC design is its final key feature, as it enables both individual differences-focused and other theoretically-valuable investigations of cognitive control function. Specifically, within the DMCC, many participants, including most of the MZ twin pairs, return for multiple waves of testing, during which the full experimental protocol is repeated. This aspect of the design provides a test of the longer-term (i.e., multiple month intervals) stability versus change present in behavioral and neural cognitive control profiles. Combined with the multi-task, multiple context nature of the design, this longitudinal data will enable richer and more rigorous investigations that can uncover latent sources of stability and change decoupled from task-specific effects and other sources of measurement error. Moreover, for the subset of DMCC participants in the HCP, there is also the possibility of relating cognitive control function with stability versus change in resting-state connectivity patterns over even longer intervals (years). Lastly, for many DMCC participants the longitudinal component is integrated with a mindfulness skills training intervention that occurs between testing waves. This component of the project provides a novel means of testing the relationship between mindfulness and cognitive control function – a relationship that has been hypothesized in many theoretical treatments (Malinowski, 2013; Y.-Y. Tang & Posner, 2009; Teper et al., 2013). Since most of the DMCC participants in the mindfulness intervention are members of MZ twin pairs (with their co-twin serving as a wait-list control), the DMCC project enables investigation of how psychologically and cognitively meaningful interventions impact relevant individual differences in cognitive control. This can be done by examining how similarity in cognitive control function among MZ twins (relative to unrelated individuals) is impacted by a discordant and randomized experimental manipulation.

We describe these key features of the DMCC project in the hopes of making other interested investigators aware of the possibilities available with the dataset. Aligned with the current push towards open science and reproducibility, we have a strong commitment to make the DMCC dataset publicly available. While the project is ongoing, we plan for a partial release by the time this paper is published (of baseline session data), and a full release after study completion. Regardless, we encourage interested investigators to contact us regarding the possibility for earlier collaboration or dataset access.

### Initial DMCC findings

To provide a first demonstration of the promise of the DMCC dataset, we conducted analyses that focused on reactive control, particularly whether there was a consistent behavioral and neural signature of this control mode. The reactive control mode has tended to receive less investigation than that of proactive control (Braver, 2012), and its characteristics of reactive control have been more difficult to describe, sometimes even confused or conflated with a reduction in proactive control. Thus, one of the design goals of the DMCC battery was to provide task conditions, across a set of well-established cognitive control paradigms, that manipulated the utilization of reactive control, distinct from proactive control. In particular, we hoped to demonstrate that utilization of the reactive control mode can be linked with enhanced cognitive control function and improved task performance (i.e., reduced interference effects). In this respect, the results clearly supported our hypotheses. In all four tasks, we observed behavioral evidence of task performance improvements under Reactive conditions, observable as reductions in interference relative to Baseline.

We found a parallel and novel pattern when examining the fMRI data, in that a set of fronto-parietal brain parcels, defined by their consistent transiently increased activation to high control demand target items during the Baseline condition, also showed a consistent *reduction* in this transient response in the Reactive condition. Three aspects of this finding are particularly noteworthy. First, because we identified the set of 34 fronto-parietal parcels based solely on their Baseline activation profiles, the activity reduction we observed in these same parcels under Reactive conditions was unbiased. Second, the set of fronto-parietal regions that we identified here are consistent with prior work, which described a similar canonical cognitive control network (Cole & Schneider, 2007; Dosenbach et al., 2008; Vincent et al., 2008). This network has also been referred to as the multiple-demand network (Camilleri et al., 2018; Duncan, 2010; Duncan & Owen, 2000), in that it exhibits consistent responsivity to increasing control demands across a range of task contexts (Assem et al., 2020; Fedorenko et al., 2013; Shashidhara et al., 2019). This set is primarily bilateral and includes not only mid-lateral prefrontal cortex and parietal cortex, but also other regions linked to cognitive control, such as the anterior insula / frontal operculum, and medial frontal cortex / dorsal anterior cingulate. Our use of a standardized atlas and parcellation scheme (Schaefer et al., 2018) facilitates identification of these regions in multiple datasets and across labs, which will promote stronger tests of cross-study consistency in anatomical localization and naming conventions. Third, because of the multi-task nature of the DMCC design, we were able to demonstrate a consistent pattern, not only for the Baseline control demand effect, but also for the Reactive activation reduction effect. This consistency increases confidence in the interpretation that the Reactive pattern is a generalizable one, occurring across different task contexts and experimental manipulations.

Indeed, reactive manipulations implemented across the four tasks were somewhat variable, and in some cases, novel. In the Stroop task, we used an item-specific proportion congruency manipulation that has been repeatedly shown in prior work (Bugg et al., 2011; Bugg & Hutchison, 2013; Gonthier, Braver, et al., 2016). In the AX-CPT, the manipulation was similar, in that we used a context-specific proportion conflict manipulation (spatial location and border color of probe items predicted the likelihood that they would be high or low conflict). To our knowledge, this is the first time that both item-specific and context-specific manipulations have been studied together in a within-subject, multi-task design. In the Sternberg task, the reactive manipulation also involved interference expectancies, but was related to familiarity, rather than congruency or conflict. Finally, in the Cued-TS paradigm, the reactive manipulation involved a punishment-related motivational context, so was quite different from the other tasks. Yet even with these heterogeneous manipulations, we observed a similar fronto-parietal neural profile for reactive control. As such, it seems warranted to refer to this fronto-parietal pattern of reduced transient activation as the *neural signature of reactive control*.

The functional relevance of this neural signature was supported by the finding of a clear brain-behavior relationship, highlighting individual differences among participants. In particular, the individuals that showed the strongest Reactive-related reduction in transient activity were the ones showing the largest behavioral benefits. The relationship was confirmed through formal statistical mediation analyses, which suggest that the impact of the Reactive experimental manipulations on task performance, were at least partially mediated by their associated impact on neural activity. Thus, we interpret the findings as indicating that the Reactive-related reduction in transient activity is functionally related (or even more provocatively, a causal factor tied) to the improvement in performance.

There were some potentially surprising aspects of these reactive control findings. First, it might be expected that an increase in cognitive control utilization would be associated with increased activation in the neural substrates that implement control functions. Yet here we found the opposite pattern, in which we attribute enhanced utilization of reactive control with a transient decrease in event-related activation. In the neuroimaging literature, decreased activity patterns co-occurring with improvements in behavioral performance markers are often referred to as reflecting enhanced efficiency in neural computation. That is, the computation may take less time, or require less resources. Indeed, efficiency-related activation patterns have been attributed to enhanced cognitive control in some prior work (Gold et al., 2013; Gray et al., 2005; Luna et al., 2010), and in particular, in item-specific proportion conditions that reflect utilization of reactive control, in tasks such as the Stroop (Blais & Bunge, 2010; Chiu et al., 2016; Grandjean et al., 2013). Thus, in some ways, the findings are consistent with the prior literature. Nevertheless, computational and mechanistic accounts of how enhanced cognitive control can give rise to reduced activation are still somewhat lacking. Thus, a fuller understanding the neural mechanisms of reactive control will still require further work, as elaborated further below.

Another potential issue related to the interpretation of the current results is that frequently, evidence of reduced activation occurs with practice or repetition of items. This effect is so common in the memory and object recognition literature that it is referred to as the repetition suppression effect (Gonsalves et al., 2005; Grill-Spector et al., 2006; Henson & Rugg, 2003; Horner & Henson, 2008; Larsson & Smith, 2011). Similarly, increasing practice or experience with task conditions is also well-established to lead to reductions in task-related activation (Chein & Schneider, 2005; Kelly & Garavan, 2004; Petersen et al., 1998). Finally, reactive control has been frequently interpreted as related to the learning of stimulus-response or stimulus-control state associations (Chiu & Egner, 2019), potentially via conflict-triggered learning mechanisms (Abrahamse et al., 2016; Verguts & Notebaert, 2008, 2009). These associations will build up with repeated exposures to stimulus trials and features. Thus, it is a possible concern that in the DMCC study design the Reactive session always occurred after the Baseline session. As a consequence, one alternative interpretation of the Reactive-related reductions in activity is that they merely reflect the increased practice that occurs with the general task conditions during the Reactive session relative to Baseline, rather than anything specific or selective regarding the engagement of reactive control. In the analyses conducted for the paper, we partially controlled for this alternative interpretation by examining reactive effects as a function of whether the Reactive conditions were performed in the second or third session (which was counterbalanced across participants). Under a pure practice or repetition interpretation of the results, both the reduction in reactive activation (and behavioral performance) would be expected to be stronger in the third session than the second. Yet no evidence of this pattern emerged. Nevertheless, stronger evidence for the selectivity and specificity of the reactive effects is still needed.

### Limitations and Future Directions

Although beyond the scope of the current paper, a key next step in the DMCC project will be to focus on determining whether a unique neural signature of proactive control can be also be identified. A central claim of the DMC framework, and a primary focus of the DMCC project was to test whether proactive and reactive control are implemented by distinct and dissociable neural mechanisms. In particular, the current findings are supportive of DMC framework tenets that reactive control is associated with transient neural mechanisms that are stimulus-triggered and linked to key features of target items (e.g., conflict, familiarity, punishment, etc). However, proactive control is postulated to involve distinct neural mechanisms that are sustained or cue-related, and which become engaged ahead of, and in anticipation, of control-demanding events. To most strongly confirm the presence of these independent neural control mechanisms, it will be critical to determine whether a double dissociation can be established, such that the Proactive condition is selectively associated with consistent changes in sustained or cue-related activity across the four tasks in the battery, while the Reactive condition is selectively associated with transient event-related effects. Moreover, establishment of such a double dissociation will be highly important for ruling out alternative interpretations of the findings, such as those related to practice or session-order confounds, such as described above.

Even without a contrasting focus on proactive control, the findings reported here can also be advanced by further explication and convergent evidence regarding underlying mechanisms. In particular, one intriguing hypothesis is that the Reactive-related reduction in transient activity may reflect more efficient signaling and propagation of control-related information among fronto-parietal regions. Furthermore, recent work has pointed to fronto-striatal pathways, particularly involving the dorsal striatum and caudate nucleus, in mediating the learning and storage of stimulus-control associations (Chiu & Egner, 2019). Thus, one important direction for future work will be complement the activation-focused analyses conducted here, with those aimed at understanding whether reactive control is also linked to systematic changes in functional or effective connectivity, among fronto-parietal and fronto-striatal regions.

Another important direction will be to exploit both the twin-based and test-retest components of the DMCC study design to further explore and support reactive (and proactive) control findings. For example, a powerful means by which to demonstrate the robustness and generalizability of the reactive control neural signature would be to test whether it is stable across repeated waves of testing. Such a finding would also be important for ameliorating concerns regarding whether the reactive control pattern is confounded with practice or repeated exposure to experimental task conditions. In other words, if the activity reductions found in the Reactive condition, relative to Baseline, were also replicated in retest sessions, this would argue against a practice-related interpretation, since in the retest sessions, all tasks would already be highly practiced and familiar. Another way to establish the functional importance of the reactive control signature would be test whether the individual difference-related brain-behavior relationships we observed were also highly similar within MZ twin pairs. If such a pattern were observed, it would clearly support the interpretation that the reactive control signature reflects a meaningful and heritable dimension of individual variation, rather than a purely state-dependent activation pattern. This type of inference would also be supported by findings that the observed reactive control patterns were linked to other sources of individual differences. For example, in prior work, we and others have suggested that reactive control might be associated with increased trait anxiety (Fales et al., 2008), risk aversion (Brown & Braver, 2007), and punishment sensitivity (Savine et al., 2010). Testing for these associations represents a rich and promising direction of exploration within the DMCC dataset, once the sample size permits such investigations to be conducted with sufficient statistical power.

Another potential limitation of the current work, and also a key direction for future analyses, relates to the constraints imposed by a univariate, as opposed to multivariate, statistical modeling approach to the DMCC data. Indeed, univariate approaches only provide information about the intensity of brain activation (i.e., increasing or decreasing), and as such are of limited utility for understanding the computations that are more likely to be encoded in activation patterns. In contrast, multivariate approaches, such as MVPA, provide richer information regarding the representational coding schemes that are implemented in different brain regions, which might be critical for disentangling and dissociating cognitive control dimensions, such as proactive and reactive. In particular, reactive and proactive control may be most strongly dissociated in terms of the neural coding of distinct control dimensions (Freund et al., 2020). Indeed, reactive control might be more strongly identified in terms of the neural coding of congruency information or stimulus-control associations, whereas proactive control is likely to be associated with the coding of task rules or goal-related information. Our group has started to explore MVPA approaches, such as classification decoding (R. Tang et al., 2020) and representational similarity analysis (RSA; Freund & Braver, 2020) with the DMCC data, and this represents a promising direction for more systematic exploration.

A related and critical question within the DMCC project is to more firmly establish the domain-generality of reactive and proactive control. Although domain-generality can explored with univariate approaches as well, such as through conjunction analysis and composite indices, as utilized in this report, such methods are less powerful, since they are susceptible to measurement error and other sources of noise. Indeed, there has been quite a bit of recent controversy and debate regarding whether univariate behavioral and neural measures of cognitive functioning are sufficiently reliable to be treated as individual difference measures and included in cross-task analyses (Bennett & Miller, 2010; Dubois & Adolphs, 2016; Elliott et al., 2020; Hedge et al., 2017; Rouder & Haaf, 2019; Whitehead et al., 2019). One solution to this problem is to use latent variable approaches, which are especially well-adapted to the multi-task, multi-condition approach utilized in the DMCC project. Because latent variable approaches examine relationships between latent factors, estimated in shared variance across tasks or conditions, they are less prone to measurement error, and as such may be more sensitive and selective in revealing the domain-generality of cognitive control dimensions. Indeed, our group has begun exploring the use of latent variable approaches to neuroimaging analyses, in large-sample datasets such as the HCP, and found them to be quite effective (Cooper et al., 2019). However, one of the main drawbacks of such approaches is that they are “data-hungry” and are likely not to be sufficiently powerful in datasets such as the DMCC until data collection is complete, with sample-sizes in the range of 200 hundred individuals.

This is again an issue for which multi-variate approaches might represent an alternative and more efficient solution (Dubois & Adolphs, 2016). A key advantage of MVPA approaches is not only that they provide richer descriptions of neuroimaging data, by more effectively pooling voxel-wise patterns, but also that they may more easily enable more abstracted measures brain activity, that are less susceptible to measurement error and artifacts (Cohen et al., 2017; Norman et al., 2006). In particular, RSA measures, which focus on the similarity in neural activation patterns, can enable examinations of second-order relationships (Haxby et al., 2014; Kragel et al., 2018; Kriegeskorte et al., 2008; Kriegeskorte & Kievit, 2013), such as the relative similarity of high control and low-control demand conditions across tasks. By focusing on cross-task and cross-session similarity structure, these types of RSA approaches are likely to be the most powerful ones for revealing whether the neural coding of reactive and proactive control does indeed show the theoretically-predicted forms of domain-generality suggested by the DMC framework.

## Conclusion

The DMCC project represents an ambitious and rigorous attempt to reveal key neural mechanisms of cognitive control. In particular, it tests the key theoretical tenets of the DMC framework, which postulate reactive and proactive modes as meaningful and distinct dimensions of cognitive control that are serve as important sources of both intra- and inter-individual variation. Towards that end we have provided initial support for these DMC tenets, by identifying a novel neural signature of reactive control, that it is: a) consistently engaged across multiple tasks; b) involves a focal set of fronto-parietal regions; c) contributes to behavioral performance enhancements observed under that conditions encourage utilization of reactive control; and d) is subject to significant individual differences. Our goal is to inform investigators interested in individual differences and cognitive control of the advantageous design features of the DMCC project, with the hope that they will be encouraged to make use of the dataset, when the data become publicly available, which we commit to doing as soon as possible. In that spirit, we hope to contribute to building a strong community of researchers working together to advance our understanding of frontal lobe function, the neural underpinnings of cognitive control, and individual differences in these domains.

## Acknowledgements

This research was supported by NIH R37 MH066078 and R21 AT009483 to T.S.B. We are grateful for the collaborative support of the Human Connectome Project, WU-Minn Consortium (Principal Investigators: David Van Essen and Kamil Ugurbil; 1U54MH091657). We thank the many individuals who contributed to the development, design and acquisition of the DMCC: Andrew Conway, Julie Bugg, Carol Cox, Marie Krug, Kevin Oksanen, Leah Newcomer, Maria Gehred, Shelly Cooper, Allison Tay, and Anxu Wang. We thank the members of the Cognitive Control and Psychopathology Lab for many fruitful discussions and feedback regarding the DMCC project.

## Notes

https://osf.io/xvzrf/

### Competing Interest Statement

The authors have declared no competing interest.

